# Comprehensive Characterization of Somatic Mutations and Gene Expression on TP53/PTEN DKO Murine Glioblastoma Cell Lines: Insights from Whole-Exome and RNA-seq Analysis

**DOI:** 10.64898/2026.11.10.765700

**Authors:** Muhammad Zain, Sumaira Nishat

## Abstract

Glioblastoma (GBM) is an aggressive brain tumor characterized by genomic instability and transcriptional plasticity. To investigate the somatic evolution of GBM, we performed an integrated analysis using Whole-Exome Sequencing (WES) and RNA-seq on a murine glioblastoma model representing two stages: early-stage transformed neural stem cells (tNSC) and late-stage malignant GBM (mGB). The comparative analysis of mutational signatures provides evidence of a two-phase evolutionary path for tNSC and mGB. Variant calling in tNSCs revealed mutations in *Pfdn2*, *Oip5*, *Trmt10a*, *Nabp2*, *Man2b1* with mGBs acquiring additional putative progression-associated genes such as *Cdk4*, *Ubap2*, *Ccdc11*, *Errfil*, and *Tead2* associated with malignancy and Temozolomide (TMZ) resistance. Samples in tNSCs exhibited signatures associated with stress-(SBS4) and development-associated (SBS40) processes. In contrast, mGB samples exhibited an increase in mutation signatures, specifically associated with replication-(SBS5) and mismatch repair (MMR) processes (SBS26). Notably, SBS26 consistently ranked in the top five mutational signatures across all samples, indicating that there may have been an early, and sustained, loss of MMR. Ti/Tv ratio also revealed elevated transversions in tNSCs supporting early instability post TP53/PTEN knockout, while mGB demonstrated normal transition-to-transversion ratio. Gene ontology reveals activities in neurodevelopmental pathways in tNSCs such as neurogenesis, axon guidance, and synapse regulation while in mGB it shifts to cellular transformation and invasion, and epidermal development. Moreover, differential expression analysis showed extreme downregulation against control neural stem cells (ctrlNSCs) indicating genomic instability and then the partial restoration of expression in mGB, resembling ctrlNSCs but with a bias toward malignant upregulations. Together this suggests, early genomic instability disrupting developmental programs and blocking differentiation, and later on replication-driven mutations and cytoskeletal remodeling reprograms the tumor into a proliferative, invasive, and therapy-resistant state.

## 1 Introduction

### 1.1 Background

Glioblastoma (GBM) is the most aggressive and lethal primary brain tumor in adults, classified as grade IV by the World Health Organization (WHO) [1]. Patients with GBM have a median survival of about 14-16 months, despite advancements in surgical resection, radiotherapy and chemotherapy [2]. The heterogeneous nature of GBM, with a combination of genetic, epigenetic, and transcriptomic alterations, leads to resistance to standard treatments and ultimately poor clinical outcomes. Thus, understanding the molecular mechanisms responsible for glioblastoma development and progression is crucial to identifying new therapeutic targets to improve patient prognosis [3].

### 1.2 Description

The evolution of glioblastoma (GBM) is believed to involve a multistep process in which the cells undergo a series of progressive events following the acquisition of somatic mutations that transform normal glial or neural stem cells into malignant glioma cells [4]. To that end, many recent publications report a number of recurrent genetic changes in GBM that are critical to GBM progression, including the mutations in *TP53, PTEN, IDH1/2, EGFR, CDK4, NF1* and *ATRX* [5]. However, *TP53* and *PTEN* are essential for the maintenance of genomic stability and for regulating signaling pathways [6]. Loss of function of *TP53* disables the DNA damage response leading to uncontrolled proliferation and accumulation of mutations, while loss of function of *PTEN*, which is an important regulator of the PI3K/AKT pathway, contributes to aberrant cell survival and metabolic reprogramming [7]. Experimental models with co-inactivation of *TP53* and *PTEN* have shown accelerated gliomagenesis and aggressive tumor characteristics, making them good systems to understand the molecular aspects of GBM development [8].

### 1.3 Scope

The availability of next-generation sequencing (NGS) has enabled global assessments of somatic mutations in the cancer genome [9]. Integrating whole-exome with RNA-seq provides a complete understanding of both somatic mutations and their functional consequences on gene expression. This allows for a more complete view of tumor initiation and progression.

This study explores molecular evolution of GBM using TP53/PTEN double knockout murine model representing pre-malignant transformed neural stem cells (tNSCs), and malignant Glioblastoma (mGBs). Somatic variants were identified by calling somatic variants and characterized using variant annotation, transition-to-transversion ratio analysis (ti/tv ratio), mutational signature analysis, and gene ontology to define the mutational processes associated with disease progression. Beyond mutation profiling, RNA-seq was integrated to evaluate transcriptomic changes accompanying tumor evolution. Differential gene expression analysis was performed to identify alterations associated with malignant transformation. Together, these complementary genomic and transcriptomic analyses enable the investigation of how genomic instability, DNA repair deficiencies, developmental dysregulation, and transcriptional reprogramming collectively contribute to the transition from transformed neural stem cells to malignant glioblastoma.

### 1.4 Objectives

In our study, we used a *TP53/PTEN* double-knockout mouse model to understand the molecular evolution of glioblastoma from pre-malignant neural stem-like cells (tNSC) to malignant glioma (mGB). We performed whole-exome sequencing (WES) to assess somatic mutation patterns, especially focusing on the Ti/Tv ratio and the spectra of high- and moderate-impact somatic mutations in important cancer-related genes. When comparing mutation patterns between tNSC and mGB samples, we sought to define the transition from early genomic instability to refined oncogenic signatures. We also assayed the biological pathways implicated with these mutations using mutational signature analysis, gene ontology enrichment, and differential gene expression analysis. Our results contribute to the mechanistic understanding of glioblastoma evolution in light of *TP53/PTEN* loss and provide therapeutic targets.

## 2 Methodology

### 2.1 Data Acquisition

Whole-exome sequencing (WES) and RNA-sequencing (RNA-seq) data used in the study were obtained from publicly accessible Gene Expression Omnibus (GEO) under the accession number GSE145559 [10]. We downloaded raw sequencing FASTQ files using the Sequence Read Archive (SRA) toolkit’s fasterq-dump. The dataset comprises *TP53/PTEN* double-knockout murine glioblastoma models, including transformed neural stem cells (tNSCs), malignant glioblastoma cells (mGBs), and control neural stem cells (ctrlNSCs) dataset which enabled a comparative analysis of somatic mutations and gene expression patterns in glioblastoma (GBM) development.

### 2.2 Variant Calling and TiTv Analysis

Variant calling was performed to identify underlying variants playing its role in GBM progression. Quality control of raw reads was performed using FastQC (v0.11.5) for both WES and RNA-seq. No trimming was applied to the exome data and was aligned to GRCm39 reference genome using `BWA-MEM` (v0.7.17). Sorting and duplicate marking were performed using `GATK` (v4.3.0.0) `MarkDuplicates`, and duplicates were removed using `Samtools` (v1.13) with the `-F 1024` filter.

Base quality score recalibration was carried out using `BaseRecalibrator` and `ApplyBQSR`. As dataset contained tumor-only samples, known polymorphic sites for recalibration were provided using the SNP reference file corresponding to the same `GRCm39` genome build, and indexed with `tabix`. Somatic variants were identified using `Mutect2`, filtered with `FilterMutectCalls` [11]. Variants obtained after filtering were annotated with Ensembl’s Variant Effect Predictor (VEP) (release 113) and downloaded annotated vcf files [12]. The transition/transversion (TiTv) ratio of the annotated variants was calculated using bcftools stats function.

### 2.3 Mutational Signature Analysis

We utilized the MutationalPatterns R package [13] for mutational signature analysis against COSMIC v3.2 data. The somatic VCF files produced after annotation were first preprocessed using the `read_vcfs_as_granges()` function, with the `Bsgenome.Mmusculus.UCSC.mm39` reference genome to ensure accurate context-dependent mutation classification. As `Bsgenome.Mmusculus.UCSC` follows UCSC-style contig naming while somatic files were aligned to NCBI’s GRCm39 reference, contig names were converted it to UCSC-style using `bcftools` to ensure compatibility during downstream analysis.

We extracted the 96 trinucleotide mutational contexts using the `mut_matrix()` function. The resulting matrix was normalized and visualized. To identify known mutational signatures, we fitted our mutation profile against the COSMIC v3.2 Single Base Substitution (SBS) signature reference set using the `fit_to_signature()` function. Cosine similarity between the observed mutation spectrum and each COSMIC signature was calculated, with a threshold of 0.9 indicating strong concordance. Signatures were normalized and then visualized using `ComplexHeatmap`.

### 2.4 GO Enrichment Analysis

To identify molecular functions and biological processes associated with somatic variants, we performed Gene Ontology (GO) enrichment analysis focusing on high- and medium-impact variants for both tNSC and mGB profiles. These were filtered based on Impact annotation provided by VEP, where variants labeled as HIGH or MODERATE impact were retained. Corresponding gene symbols were extracted and de-duplicated for downstream analysis. GO enrichment analysis was performed using `clusterProfiler` (4.16.0) [11]. Gene list was converted to Entrez gene ID using `bitr()` function with the `org.Mm.eg.db` annotation package.

The enrichGO() function was used to identify overexpressed GO to identify biological process (BP), molecular function (MF), and cellular components (CC) categories. Multiple testing correction was applied using the Benjamini–Hochberg false discovery rate (FDR) method and adjusted p-values (q-values) < 0.05 were considered significantly enriched. The top enriched terms were visualized using dotplot and emapplot.

### 2.5 Differential Expression Analysis

RNA-seq samples underwent quality control and head trimming using `Trimmomatic` (v0.39). The reference genome was indexed using `HISAT2` (v2.2.1), and raw counts were obtained using `featureCounts` (v1.6.0). Differential expression analysis was conducted using `DESeq2` [14], comparing control NSC (ctrlNSC), transitional NSC (tNSC), and malignant GBM (mGB) samples. Variance stabilizing transformation (VST) was applied for normalization. Differentially expressed genes (DEGs) were identified using an adjusted p-value threshold of 0.05 and a log2 fold change of 1. Heatmaps were generated using `ComplexHeatmap`, and volcano plots were constructed using `ggplot2`.

## 3 Results

### 3.1 Whole-Exome Analysis

The GBM dataset (GSE145559) offers a comprehensive characterization of GBM progression at malignant and pre-malignant stages. It comprises whole-exome sequencing (WES) and RNA-seq data of neural stem cells (NSCs) isolated from double knock-out *pten/tp53* mice at distinct progression stages. The earliest stage, tNSC0 was collected two weeks after tamoxifen-induced *pten/tp53* deletion. At this stage, the expansion of rostral migratory stream (RMS) was observed suggesting early tumorigenic potential. Tumor cells progressed to established GBM with mGB0 representing high-grade glioma emerging 12 months post NSC expansion. Tumor-derived cell lines (tNSC1-tNSC3, mGB1-mGB2) were established though serial transplantation into immunocompetent C57/Bl6N mice, exhibiting 100% penetrance and reduced survival time. The dataset provides an in-depth investigation of molecular progression of GBM at pre-malignant and fully malignant stages [10].

#### 3.1.1 Variant Calling and Annotation Reveal Distinct Evolution between Pre-Malignant and Malignant Profiles

To investigate the molecular basis of glioblastoma (GBM) progression, variant calling revealed a significant increase in variant burden in mGB samples compared to their tNSC counterparts. Specifically, mGB samples exhibited a median of 437,425 variants, whereas tNSC sample contained a median of 201,179 variants (Table 1). This implies a large burden of mutations that occur during the transition from pre-malignant neural stem cells to fully transformed GBM cells.

**Table 1:** Summary of Variants across samples.

| **Category** | **tNSC0** | **tNSC1** | **tNSC2** | **tNSC3** | **mGB0** | **mGB1** | **mGB2** |
|---|---|---|---|---|---|---|---|
| Total Variants Processed | 233,589 | 190,350 | 199,085 | 181,694 | 475,702 | 424,497 | 412,076 |
| Novel Variants | 70,414 | 41,245 | 65,185 | 51,624 | 51,028 | 42,546 | 46,488 |
| Existing Variants | 163,175 | 149,105 | 133,900 | 130,067 | 424,674 | 381,951 | 365,588 |

**Table 2:** Sample-based High impact variants with variant allele frequency in tNSCs.

| Sample | Gene | CHR:POS | Consequence(s) | VAF (AD/DP) | Existing/Novel | VEP Imp. |
|---|---|---|---|---|---|---|
| **tNSC0** | *Pfdn2* | NC_000067.7:171184184–171184270 | Splice_donor_variant, splice_donor_5th_base_variant, intron_variant | 189/741=0.2551 | rs3388516203 | High |
|  | *Oip5* | NC_000068.8:119448705–119448731 | Splice_donor_variant, frameshift_variant | 53/168=0.3155 | rs3391829705 | High |
|  | *Creb3l2* | NC_000072.7:37313250 | splice_acceptor_variant, non_coding_transcript | 58/216=0.2685 | rs3413082358 | High |
|  | *Rpl8* | NC_000081.7:76788648–76788736 | Splice_donor_variant, coding_seq_variant | 68/614=0.1107 | rs3406166997 | High |
| **tNSC1** | *Pfdn2* | NC_000067.7:171184184–171184270 | splice_donor_variant, intron_variant | 180/873=0.2062 | rs3388516203 | High |
|  | *Oip5* | NC_000068.8:119448705–119448731 | frameshift_variant | 44/156=0.2821 | rs3391829705 | High |
|  | *Trmt10a* | NC_000069.7:137860559 | Frameshift_variant, splice_region_variant | 91/433=0.2102 | Novel | High |
|  | *Man2b1* | NC_000074.7:85817900 | splice_acceptor_variant | 59/151=0.3907 | Novel | High |
|  | *Huwe1* | NC_000086.8:150623084–150623085 | frameshift_variant | 38/111=0.3423 | Novel | High |
| **tNSC2** | *Pfdn2* | NC_000067.7:171184184–171184270 | splice_donor_variant, intron_variant | 159/735=0.2163 | rs3388516203 | High |
|  | *Trmt10a* | NC_000069.7:137860515–137860516 | frameshift_variant | 353/883=0.3998 | Novel | High |
|  | *Trmt10a* | NC_000069.7:137860559 | Frameshift_variant, splice_region_variant | 93/508=0.1831 | Novel | High |
|  | *Fam3c* | NC_000072.7:22328658–22328659 | Frameshift_variant, NMD_transcript | 49/142=0.3451 | Novel | High |
|  | *Man2b1* | NC_000074.7:85817900 | splice_acceptor_variant | 55/161=0.3416 | Novel | High |
|  | *Nabp2* | NC_000076.7:128244086–128244181 | splice_acceptor_variant, intron_variant | 205/762=0.269 | rs3401780928 | High |
|  | *Kdm1b* | NC_000079.7:4723406–47234065 | splice_donor_variant, non_coding_transcript | 129/345=0.3739 | rs3404141580 | High |
|  | *Trem1* | NC_000083.7:48544384 | splice_donor_variant | 21/51=0.412 | Novel | High |
| **tNSC3** | *Pfdn2* | NC_000067.7:171184184–171184270 | splice_donor_variant, intron_variant | 132/662=0.1994 | rs3388516203 | High |
|  | *Trmt10a* | NC_000069.7:137860515–137860516 | frameshift_variant | 345/871=0.3961 | Novel | High |
|  | *Trmt10a* | NC_000069.7:137860559 | Frameshift_variant, splice_region_variant | 56/500=0.112 | Novel | High |
|  | *Fam3c* | NC_000072.7:22328658–22328659 | Frameshift_variant, NMD_transcript | 35/124=0.2823 | Novel | High |
|  | *Man2b1* | NC_000074.7:85817900 | splice_acceptor_variant | 48/167=0.2874 | Novel | High |
|  | *Nabp2* | NC_000076.7:128244086–128244181 | splice_acceptor_variant, intron_variant | 206/727=0.2834 | rs3401780928 | High |
|  | *Trem1* | NC_000083.7:48544384 | splice_donor_variant | 12/38=0.3158 | Novel | High |

**Table 3:** Sample-based High impact variants with variant allele frequency in tNSCs.

| **Sample** | **Gene** | **CHR:POS** | **Consequence(s)** | **VAF (AD/DP)** | **Existing/Novel** | **VEP Imp.** |
|---|---|---|---|---|---|---|
| **mGB0** | *Pfdn2* | NC_000067.7:171184184–171184270 | Splice_donor_variant, splice_donor_5th_base_variant, coding_seq_variant, intron_variant | 212/887=0.239 | rs3388516203 | High |
|  | *Trmt10a* | NC_000069.7:137860559–137860559 | Frameshift_variant, splice_region_variant | 92/490=0.1878 | Novel | High |
|  | *Itga6* | NC_000068.8:71668773–71668773 | splice_donor_variant | 36/136=0.2647 | Novel | High |
|  | *Itga6* | NC_000068.8:71671021–71671129 | Splice_acceptor_variant, splice_donor_variant, splice_donor_5th_base_variant, splice_poly-pyrimidine_tract_variant, intron_variant | 100/616=0.1623 | rs3391484658 | High |
|  | *Itga6* | NC_000068.8:71671319–71671410 | Splice_acceptor_variant, splice_donor_variant, splice_donor_5th_base_variant, splice_poly-pyrimidine_tract_variant, intron_variant | 115/676=0.1701 | rs3391738268 | High |
|  | *Itga6* | NC_000068.8:71671494–71671592 | Splice_acceptor_variant, splice_donor_variant, splice_donor_5th_base_variant, splice_poly-pyrimidine_tract_variant, intron_variant | 107/828=0.1292 | rs3391764905 | High |
|  | *Ubap2* | NC_000070.7:41195525–41195607 | Splice_acceptor_variant, splice_donor_variant, splice_donor_5th_base_variant, splice_poly-pyrimidine_tract_variant, intron_variant | 96/618=0.1553 | rs3394117845 | High |
|  | *Ubap2* | NC_000070.7:41195671–41195744 | Splice_acceptor_variant, splice_donor_variant, splice_donor_5th_base_variant, coding_seq_variant, intron_variant | 110/697=0.1578 | rs3507731576 | High |
|  | *Ubap2* | NC_000070.7:41195851–41195941 | Splice_acceptor_variant, splice_donor_variant, splice_donor_5th_base_variant, coding_seq_variant, intron_variant | 165/754=0.2188 | rs3394207560 | High |
|  | *Ubap2* | NC_000070.7:41199584–41199714 | Splice_acceptor_variant, splice_donor_variant, splice_donor_5th_base_variant, coding_seq_variant, intron_variant | 90/427=0.2108 | rs3393882192 | High |
|  | *Nabp2* | NC_000070.7:41199584–41199714 | Splice_acceptor_variant, splice_donor_variant, splice_donor_5th_base_variant, coding_seq_variant, intron_variant | 200/873=0.2291 | rs3393882192 | High |
|  | *CDK4* | NC_000070.7:41199584–41199714 | Splice_acceptor_variant, splice_donor_variant, splice_donor_5th_base_variant, coding_seq_variant, intron_variant | 128/1106=0.1157 | rs3393882192 | High |
|  | *CDK4* | NC_000076.7:126901872–126901969 | Splice_acceptor_variant, splice_donor_variant, splice_donor_5th_base_variant, splice_poly-pyrimidine_tract_variant, intron_variant | 177/1195=0.1481 | rs3401770155 | High |
|  | *Ywhaq* | NC_000078.7:21441793–21441793 | frameshift_variant | 53/169=0.3136 | Novel | High |
|  | *Cnih1* | NC_000080.7:47021637–47021637 | Frameshift_variant, splice_region_variant | 88/510=0.1725 | Novel | High |
|  | *Apobec2* | NC_000083.7:48730177–48730177 | stop_gained, frameshift_variant | 36/262=0.1374 | Novel | High |
|  | *Ccdc117* | NC_000077.7:5481533–5481533 | stop_gained, frameshift_variant, splice_region_variant | 65/238=0.2731 | Novel | High |
| **mGB1** | *Pfdn2* | NC_000067.7:171184184–171184270 | Splice_donor_variant, splice_donor_5th_base_variant, coding_seq_variant, intron_variant | 195/821=0.2375 | rs3388516203 | High |
|  | *Ubap2* | NC_000070.7:41195525–41195607 | Splice_acceptor_variant, splice_donor_variant, splice_donor_5th_base_variant, splice_poly-pyrimidine_tract_variant, intron_variant | 110/655=0.1669 | rs3394117845 | High |
|  | *Ubap2* | NC_000070.7:41195671–41195744 | Splice_acceptor_variant, splice_donor_variant, splice_donor_5th_base_variant, coding_seq_variant, intron_variant | 127/702=0.1809 | rs3507731576 | High |
|  | *Ubap2* | NC_000070.7:41195851–41195941 | Splice_acceptor_variant, splice_donor_variant, splice_donor_5th_base_variant, coding_seq_variant, intron_variant | 210/851=0.2467 | rs3394207560 | High |
|  | *Nabp2* | NC_000070.7:41199584–41199714 | Splice_acceptor_variant, splice_donor_variant, splice_donor_5th_base_variant, coding_seq_variant, intron_variant | 100/434=0.2304 | rs3393882192 | High |
|  | *Errfi1* | NC_000070.7:150949726–150949820 | Splice_acceptor_variant, splice_donor_variant, splice_donor_5th_base_variant, splice_poly-pyrimidine_tract_variant, intron_variant | 159/901=0.1765 | Novel | High |
|  | *Tead2* | NC_000073.7:44869967–44870064 | Splice_donor_variant, splice_donor_5th_base_variant, coding_seq_variant, intron_variant | 62/513=0.1209 | rs3397829598 | High |
|  | *CDK4* | NC_000076.7:126901872–126901969 | Splice_acceptor_variant, splice_donor_variant, splice_donor_5th_base_variant, splice_poly-pyrimidine_tract_variant, intron_variant | 105/803=0.1308 | rs3401770155 | High |
|  | *Nabp2* | NC_000076.7:128244086–128244181 | Splice_acceptor_variant, splice_donor_variant, splice_donor_5th_base_variant, coding_seq_variant, intron_variant | 202/717=0.3274 | Novel | High |
|  | *Cnih1* | NC_000080.7:47021637–47021637 | Frameshift_variant, splice_region_variant | 124/479=0.2589 | Novel | High |
| **mGB2** | *Pfdn2* | NC_000067.7:171184184–171184270 | Splice_donor_variant, splice_donor_5th_base_variant, coding_seq_variant, intron_variant | 207/1182=0.1751 | rs3388516203 | High |
|  | *Ubap2* | NC_000070.7:41195525–41195607 | Splice_acceptor_variant, splice_donor_variant, splice_donor_5th_base_variant, splice_poly-pyrimidine_tract_variant, intron_variant | 214/816=0.2623 | rs3394117845 | High |
|  | *Ubap2* | NC_000070.7:41195671–41195744 | Splice_acceptor_variant, splice_donor_variant, splice_donor_5th_base_variant, coding_seq_variant, intron_variant | 172/852=0.2019 | rs3507731576 | High |
|  | *Ubap2* | NC_000070.7:41195851–41195941 | Splice_acceptor_variant, splice_donor_variant, splice_donor_5th_base_variant, coding_seq_variant, intron_variant | 275/1000=0.277 | rs3394207560 | High |
|  | *Errfi1* | NC_000070.7:150949726–150949820 | Splice_acceptor_variant, splice_donor_variant, splice_donor_5th_base_variant, splice_poly-pyrimidine_tract_variant, intron_variant | 173/916=0.1889 | Novel | High |
|  | *Tead2* | NC_000073.7:44869967–44870064 | Splice_donor_variant, splice_donor_5th_base_variant | 107/817=0.131 | rs3397829598 | High |
|  | *Nabp2* | NC_000076.7:128244086–128244181 | splice_acceptor_variant, splice_donor_5th_base_variant, coding_seq_variant, intron_variant | 245/1077=0.2275 | Novel | High |
|  | *Cdk4* | NC_000076.7:126901872–126901969 | splice_acceptor_variant, splice_donor_variant, splice_donor_5th_base_variant, splice_poly-pyrimidine_tract_variant, intron_variant | 105/803=0.1308 | rs3401770155 | High |
|  | *Cnih1* | NC_000080.7:47021637–47021637 | frameshift_variant, splice_region_variant | 175/616=0.2841 | Novel | High |

The function distribution of variant types offered meaningful patterns across groups. Missense mutations represented 30-32% of the total variants in mGB samples versus 25-31% in tNSC samples. In contrast, we found no substantial frameshift mutations, in-frame indels, or stop-gained or -lost variants, which were absent or rare in both group.

#### 3.1.2 Impact-Based Classification Highlights Early and Late Oncogenic Events

Impact based variants revealed coherent and recurrent hits across the samples. In tNSC and mGB samples, *Pfdn2* (rs3388516203) was present in all samples with relevant variant allele frequency range in 0.1994–0.2551. *Pfdn2* promotes the assembly of actin and tubulin to their corresponding polymers. This plays an oncogenic role in GBM, breast, pancreatic, and colon cancers when overexpressed [15]. Opa interacting protein 5 (*Oip5*) was present in tNSC0 and tNSC1 (VAF 0.3155 and 0.2821), which stabilizes the *E2F1* that controls the cell proliferation and DNA replication by controlling G1 to S phase transition [16]. *Trmt10a* was also present in tNSC1, tNSC2, and tNSC3 as Novel variants VAF 0.1835–0.3861, which encodes a tRNA methyltransferase responsible for m^1^G9 modification, a critical step for tRNA stability and translational fidelity [17]. The progression of *Trmt10a* in later samples implies positive selection for clones that have compromised RNA modification machinery, in result enhancing tumorigenic potential.

Several additional, stage-associated high-impact events were also identified. Recurrent splice-acceptor variants in *Man2b1* were present from tNSC1 onward (VAF 0.29–0.39), implicating altered lysosomal glycoprotein processing in the evolving clones [18]. A frameshift/NMD-predicted variant in *Fam3c* appeared in tNSC2 and persisted into tNSC3 (VAFs 0.28–0.35), consistent with acquisition of factors associated with invasion and phenotypic plasticity [19]. Mutations affecting genome maintenance and chromatin regulation were detected as well: *Huwe1* frameshift (tNSC1, VAF 0.39), *Nabp2* splice variants (tNSC2–tNSC3, VAF 0.27–0.29), and *Kdm1b* splice donor variant (tNSC2, VAF 0.37). Finally, *Trem1* splice-donor variants were observed in tNSC2 and tNSC3 (VAFs 0.3158–0.412), suggesting that immune-related signaling pathways may be perturbed during later stages of clonal evolution [20].

Taken together, the variant distribution across samples defines a clear trajectory: early events include recurrent disruption of *Pfdn2* together with *Oip5* -associated alterations that are compatible with cytoskeletal remodeling and increased proliferation; subsequent acquisition and expansion of *Trmt10a* and other novel high-impact variants mark a transition toward altered RNA processing, metabolic/lysosomal remodeling, genome instability, and immune signaling perturbation. The VAF dynamics particularly the rise of specific *Trmt10a* alleles support a model of positive selection for clones bearing defects in translational regulation, which may contribute to progressive tumorigenic potential in this system.

Multiple high-impact variants were identified in mGB samples, several of which were recurrent. A splice donor variant in *Pfdn2* (rs3388516203) was consistently present in all three mGB datasets (mGB0, mGB1, and mGB2), with variant allele frequencies (VAFs) between 0.17–0.24. Similarly, splice region variants in *Ubap2* were detected at different genomic positions, showing VAFs of 0.15–0.28, indicating recurrence across samples. *Ubap2* acts as an oncogene by targeting miR-1205 and miR-382 leading to *GPRC5A* [21].

Recurrent alterations were also found in *Nabp2* (rs3393882192) and *Cdk4* (rs3401770235), observed in more than one dataset with VAFs ranging from 0.11–0.23. *Cdk4* is a known oncogene that contributes to tumor development and resistance to therapies such as Temozolomide [22]. Additional events were noted in *Tead2* and *Errfi1*, though these were not shared by all replicates.

Several novel variants of high functional impact were detected. In mGB0, *Trmt10a* carried both a frameshift and splice region alteration (VAF 0.18), while *Apobec2* harbored a stop-gained mutation (VAF 0.14). A frameshift mutation in *Ywhaq* (VAF 0.31) was detected, and *Cnih1* exhibited recurrent frameshift changes across all three samples (VAFs 0.17–0.28). A unique stop-gained variant was identified in *Ccdc117* (VAF 0.27) in mGB0.

Taken together, the mGB cohort demonstrated recurrent disruptions affecting splice regulation (e.g., *Pfdn2*, *Ubap2*, *Nabp2*, *Itga6*) alongside novel truncating or frameshift variants in *Trmt10a*, *Apobec2*, *Ywhaq*, and *Cnih1*.

#### 3.1.3 Transition-to-Transversion (Ti/Tv) Ratios Reveal Evolutionary Shifts in Mutational Mechanisms

The transition-to-transversion (Ti/Tv) ratio is an established marker for the nature of mutational processes at work and their stability in genomics. In cancer studies, Ti/Tv ratios offer evidence of influences on DNA stability and mutation accumulation during tumorigenesis. To investigate the mutational trajectories involved in glioblastoma evolution, we calculated Ti/Tv ratios using the somatic variants found in tNSC’s and mGB’s.

The tNSC samples demonstrated a significant shift towards transversions, as seen in their comparatively lower Ti/Tv ratios; tNSC0 (1.26), tNSC1 (1.66), tNSC2 (1.18), and tNSC3 (1.40). In somatically occurring cells, expected Ti/Tv ratios are 2.0 - 2.1, thus, the observed ratios imply elevated genomic instability and breakdown in proper DNA repair fidelity [23]. An increase in transversions is often associated with mutagenic stressors (external or internal) rather than a replication error [24]. In our model, the reasons are likely caused by the loss of TP53 and PTEN, two tumor suppressors that are instrumental in maintaining the genomes integrity. TP53 loss affects DNA damage response and PTEN loss has been associated with oxidative stress and sustained cell proliferation [25]. Collectively, these molecular perturbations created a genomic environment that biases toward error-prone repair pathways that increase transversion events.

In contrast, the mGB samples (mGB0, mGB1, mGB2) consistently had higher Ti/Tv ratio (1.96, 2.02, 2.00, respectively), which indicate relative enrichment of transition events that are seen in malignant glioblastoma. This pattern suggests a transition away from the wider mutation space seen in tNSCs under stress to a narrower mutation context, regulated by endogenous regulatory mechanisms and evolutionary selection. Transition mutations, particularly C to T and G to A, often result from spontaneous or replication-associated deamination of methylated cytosines, both of which are more prevalent during extended periods of cellular replication and prior to stabilization of the cells. The increased transition bias may be indicative of APOBEC-mediated mutagenesis, which is associated with clustering of C to T mutations in specific sequence contexts and has been implicated in multiple forms of cancer [26].

The difference in Ti/Tv profiles between tNSC and mGB tumors emphasizes an inherent change in mutation processes that occurs during glioblastoma evolution. The mutation spectrum in early premalignant tNSCs with Ti mutations is mainly made up of transversions, a sign of mutational stress likely due to oxidative damage, evasive surveillance, and poor fidelity of the repair pathways. As the tumors transition from premalignant to an mGB stage, the mutation spectrum changes to transitions. This scenario can be seen in established malignancies, where endogenous processes and selective pressures constrain the fixation of driver mutations. In summary, this Ti/Tv analysis points to a significant transition in the mutational landscape of glioblastoma: from an unstable and transversion-rich mutation accrual in early transformation, to a stabilized transition-rich mutation profile in advanced malignancy. We conclude that the change from Ti/Tv patterns not only reflects changes in the biology of the tumor cells, but is likely also representative of the changing biological responses to their own intrinsic damage and the selective pressures they encountered as they progressed through time. This study suggests that the Ti/Tv ratio could be a practical surrogate for evaluating the mutational environment and tumor development stages in glioblastoma models.

#### 3.1.4 Mutational Signature Analysis Reveals Divergent Mutagenic Processes in tNSC and mGB Samples

We assessed single-base substitution (SBS) mutational signature profiling using COSMIC v3.2 [27] for each sample (tNSC0-3 and mGB0-2) in order to elucidate the underlying mutagenic processes. The analysis resulted in both overlapping and distinct mutational processes of the transformed neural stem cells (tNSCs) and the murine glioblastoma (mGB) samples.

All samples included a common core of mutational signatures, with SBS12, SBS26, SBS37, and SBS89 accounting for a majority of signatures. Importantly, SBS26, a signature associated with mismatch repair (MMR) deficiency [28] was repeatedly identified among the five largest contributing signatures in all tNSC and mGB samples demonstrating the existence of genomic instability. This MMR-deficient mutational process was potentially present throughout both sample contexts.

However, the groups also had key differences. All mGB samples uniquely exhibited very high contributions from SBS5, a clock-like signature associated with replicative aging and commonly found in proliferative tissues [29]. However, in all tNSCs, SBS5 was absent or low in number and there was instead enrichment for SBS40, another clock-like signature that is age-related but considered somewhat heterogeneous in its origins. In both cases, its presence in tNSCs and absence in mGBs suggests that tNSCs retain their mutational history of a developmental or differentiation history, while mGBs accumulate more replication-related damage.

Interestingly, SBS4, usually arising from tobacco-based carcinogens and from exposure to benzo[a]pyrene, was found in the top contributors in several of the tNSC samples (tNSC0, tNSC2, tNSC3) and was absent in all mGBs. Although exposure to tobacco-associated mutagens is not tenable in these models, this may be reflective of culture-associated oxidative stress, environmental contamination, or an unknown endogenous process mimicking this signature. Furthermore, SBS36, also linked to MMR deficiency, appeared only in one tNSC sample (tNSC2), possibly indicating a sample-specific hypermutator phenotype. The persistence of SBS12, SBS37, and SBS89 in nearly all samples, despite their unknown etiologies suggests a background mutational process shared by both cell types, possibly related to their murine origin or common experimental conditions.

These results collectively indicate that mGB samples exhibit a higher burden of replication driven mutations, reflected by SBS5 and SBS37, and may experience more proliferative stress. In contrast, tNSCs are characterized by signatures linked to early developmental history or culture-induced stress, such as SBS4 and SBS40. The shared enrichment of SBS26 points to MMR deficiency as a key underlying mechanism of genomic instability in both groups.

#### 3.1.5 Gene Ontology Enrichment Analysis Reveals Stage-Specific Biological Processes

In order to identify the functional implications of somatic mutations in both pre-malignant (tNSC) and malignant (mGB) glioblastoma stages, we conducted Gene Ontology (GO) enrichment analysis on high-and moderate-impact variants. We observed marked differences in enriched biological processes between the two groups, indicating the unique evolutionary paths of the two groups (Figure 4 and 5).

**Figure 1:**
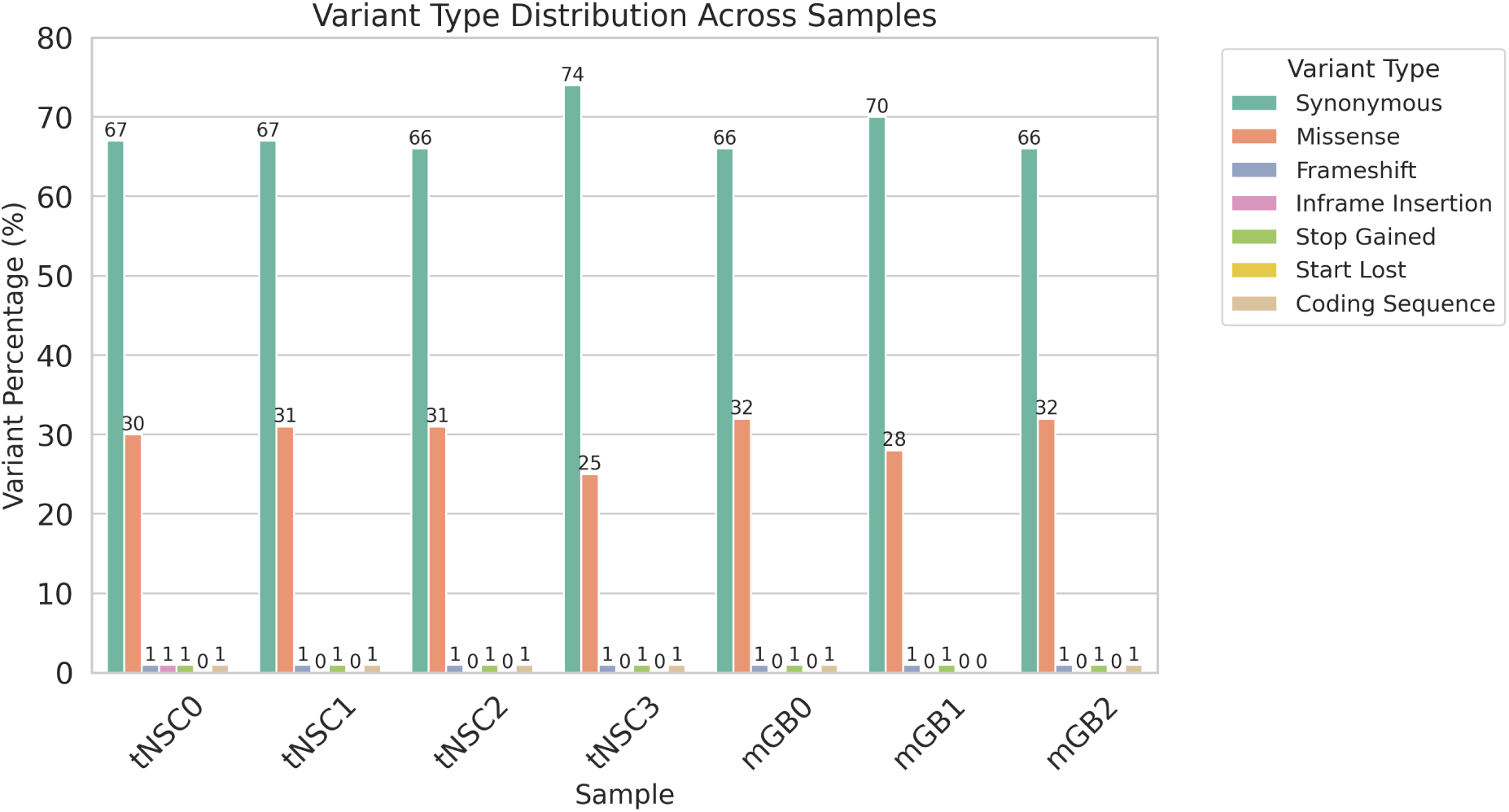
Variant type distribution across tNSC and mGB. The figure illustrates the comparative landscape of mutation types (e.g., SNPs, insertions, deletions) between pre-malignant transformed neural stem cells and malignant glioblastoma samples.

**Figure 2:**
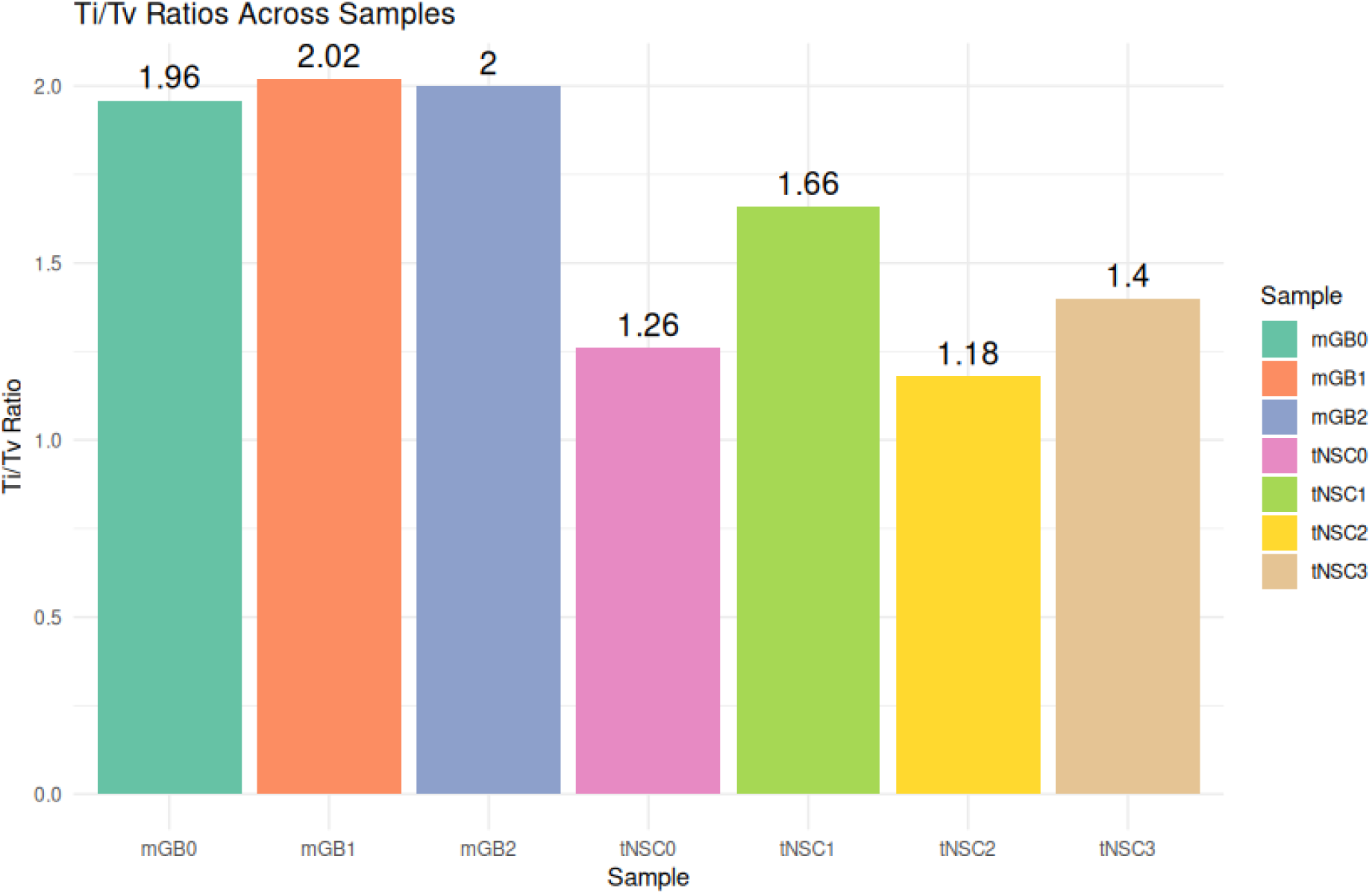
Transition/Transversion ratio across samples. The figure illustrates the comparative landscape of Ti/Tv across all the samples.

**Figure 3:**
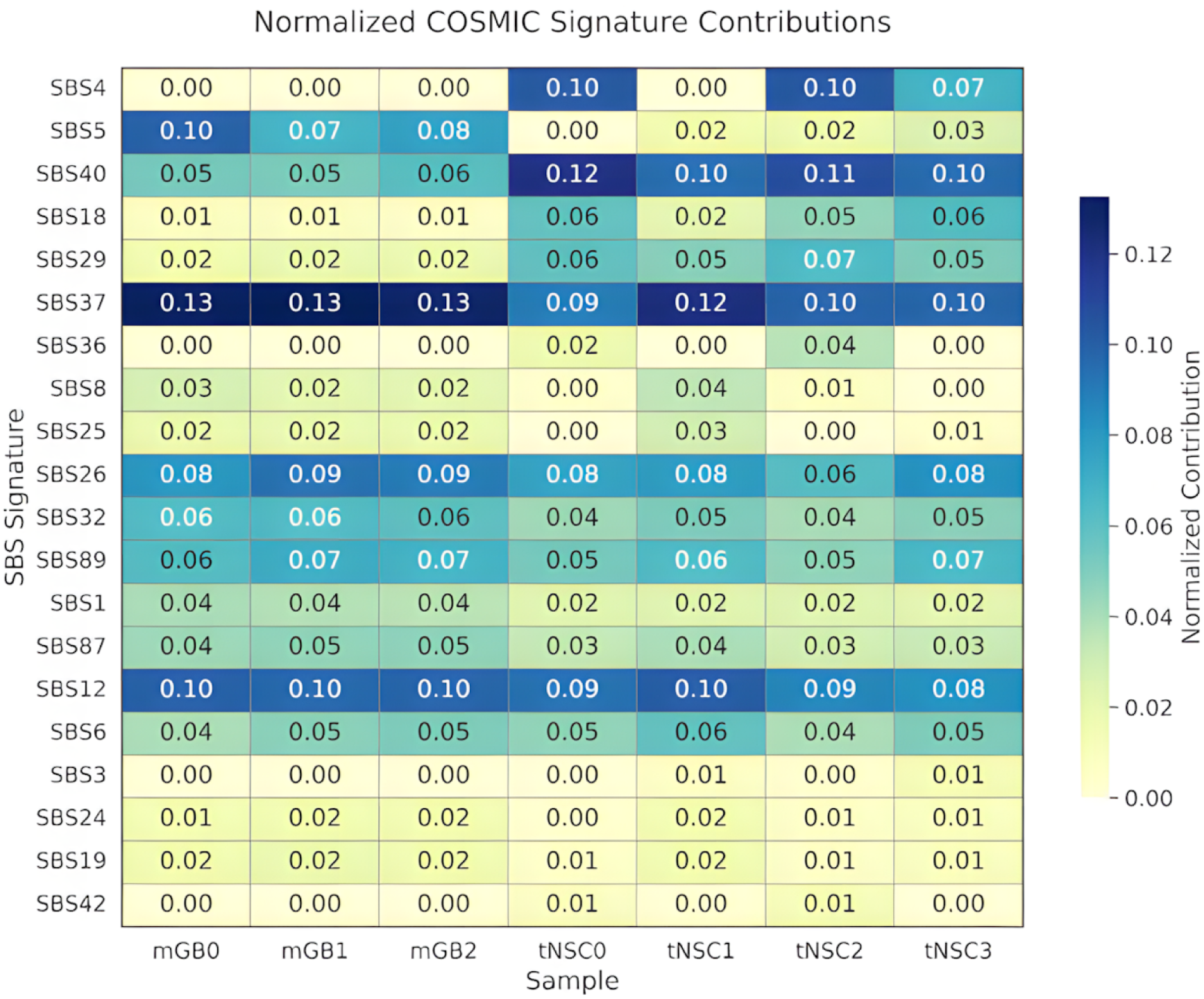
SBS profile of mGB and tNSC samples.

**Figure 4:**
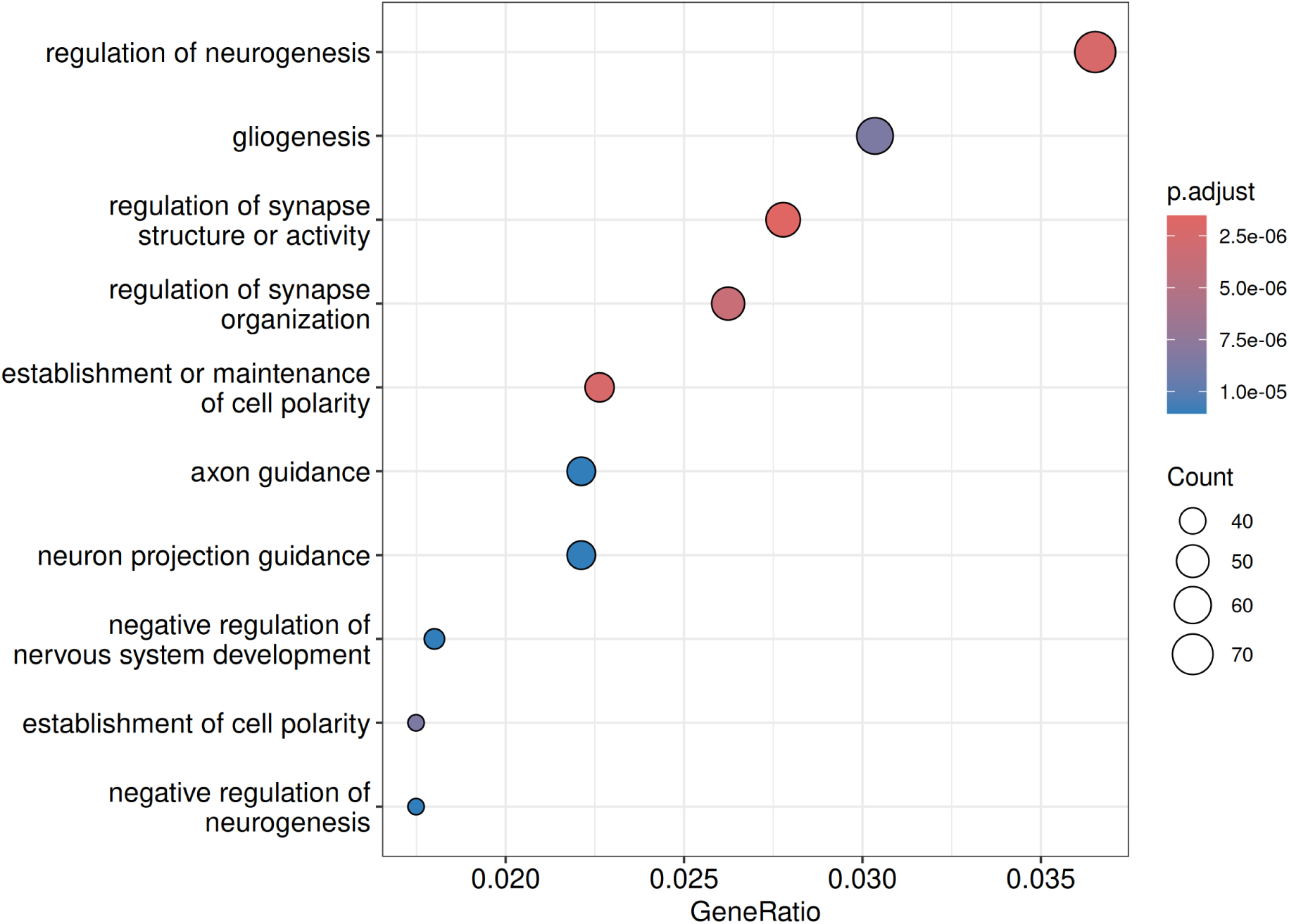
Gene Ontology of tNSC.

**Figure 5:**
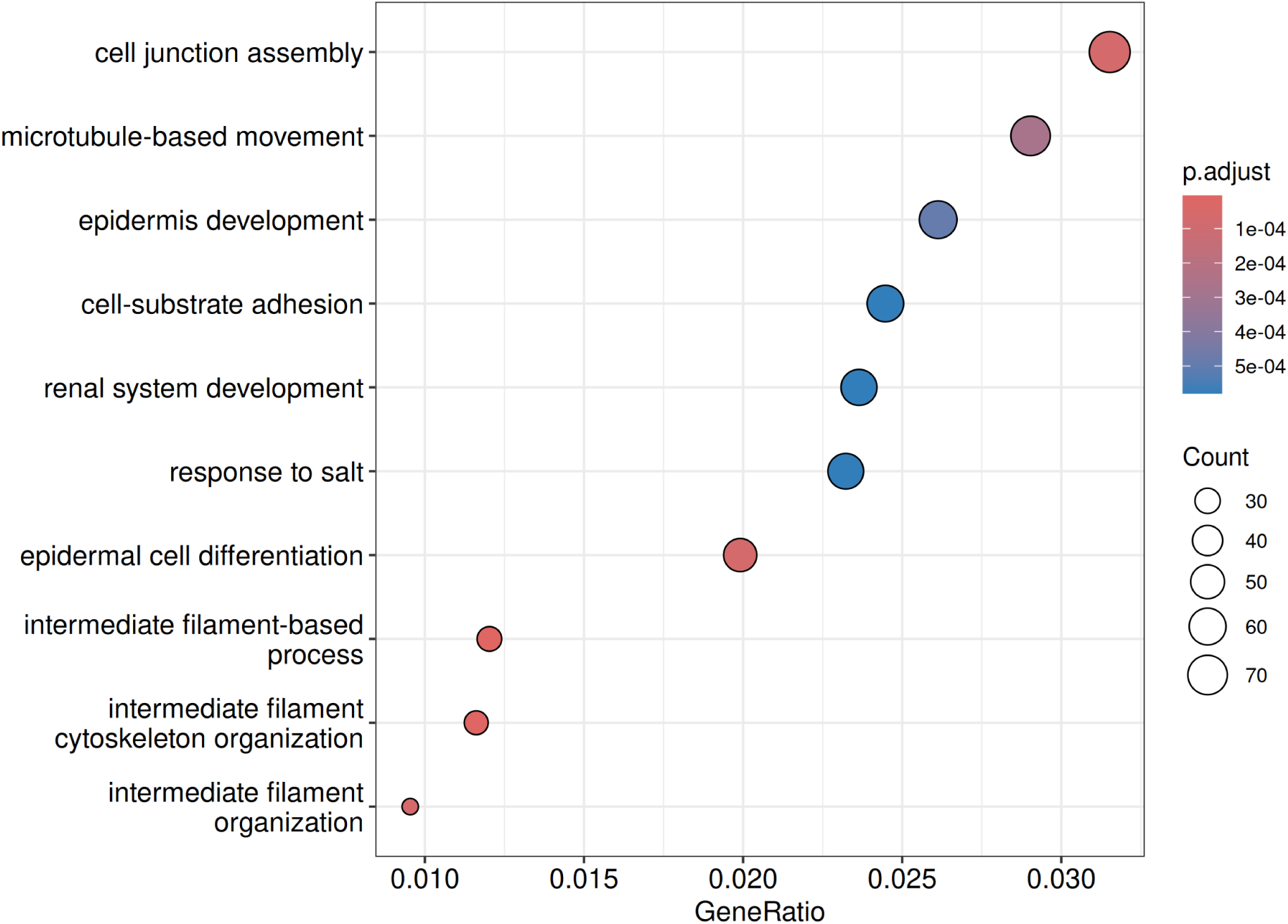
Gene Ontology of mGB.

**Figure 6:**
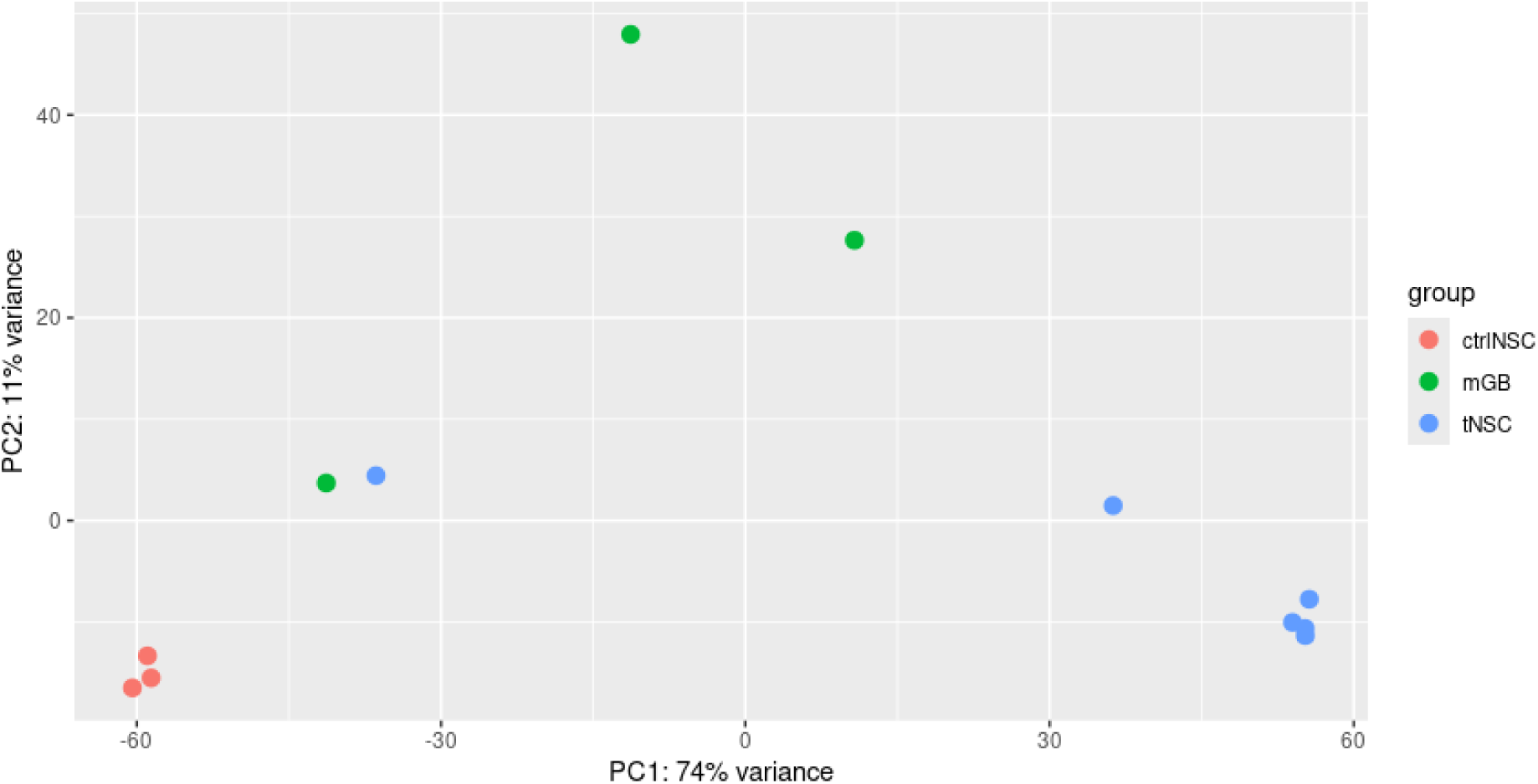
Principle Component Analysis across all profiles.

GO terms in tNSC samples were associated with neurodevelopmental processes, such as, regulation of neurogenesis (p.adjust = 2.5e-06), gliogenesis (p.adjust = 5.0e-06), and axon guidance (p.adjust = 1.0e-05) (Figure 4 (a)). These terms are consistent with their neural stem-like origins and highlight the early disturbances of pathways that are essential for differentiation [30]. Of note, regulation of synapse structure or activity (p.adjust = 7.5e-06), and negative regulation of neurogenesis (p.adjust = 0.035) are terms that support a dual role of somatic mutations, with some leading to the disallowed proliferation and some inhibiting normal differentiation. The enrichment of these terms is reconciled by the predominance of transition/transversion mutational processes (Ti/Tv = 1.18–1.66) and genomic instability associated with TP53/PTEN loss, described in Section 3.1.3.

In comparison mGB samples were enriched for terms associated with the cellular transformation supported by cytoskeletal remodeling and microenvironment interaction, which included cell junction assembly (p.adjust = 1e-04), microtubule based movement (p.adjust = 2e-04), and intermediate filament organization (p.adjust = 5e-04). These terms are significant for the acquisition of invasive and proliferative phenotypes that define advanced glioblastoma. For example, microtubule based movement supports recurrent mutations in LRP1B from mGB samples that correlate with higher glioma cell migration, a driver of tumor cell migration and treatment resistance. Intermediate filament cytoskeleton organization was also identified, possibly through glial marker expression (e.g.. GFAP) which is a responsible for the malignant behavior of gliomas. The enrichment for terms such as epidermis development (p.adjust= 3e-04) may be less typical for brain tumors, especially in the context of epithelial-like plasticity associated with mesenchymal transition in aggressive glioblastoma subtypes.

### 3.2 RNA-seq

#### 3.2.1 Principal Component Analysis Differentiates Sample Groups Based on Expression Variance

PCA was used in conjunction with hierarchical clustering in order to investigate the dominant axes of variance present in the transcriptomic data [31]. PCA1 explained 74% of the total variance in the dataset and was able to separate the ctrlNSC (left) from the tNSC and the mGB (upper) groups. Next, PC2 explained an additional 11% of the variance and was able to primarily separate the tNSC samples from the mGB samples, confirming their separate transcriptional states.

The ctrlNSC samples were clustered tightly together indicating similar transcriptional states. The tNSC samples were more dispersed than the ctrlNSC samples indicating intra-group variance and possibly a greater heterogeneity due to variability in tumor exposure, longer timescale of sampling, and epigenetic reprogramming, amongst other factors. Notably, tNSC2 and tNSC1 are closer to the mGB cluster indicating they might be candidates for experiencing early-stage tumorigenic transformation. In contrast, tNSC3 samples are clustered near the ctrlNSC samples, which may indicate they still contain some stem-like properties or may be transformed at a later time point.

Taken together, our data indicate that there is a marked distinction in transcriptional profiles and states that distinguishes normal from tumorigenic states, with tNSC representing an intermediate and transitional space in the PCA space.

#### 3.2.2 Transcriptomic Landscape Reveals Distinct Clustering Between tNSC and mGB

To characterize the transcriptional dynamics across experimental conditions, we first performed unsupervised hierarchical clustering and principal component analyses (PCA) on all gene expression profiles [32]. The global heatmap (Figure 7) presents a striking separation of samples into tNSC (transformed neural stem cell), mGB (malignant glioblastoma), and ctrlNSC (control neural stem cell) corresponding to three major groupings. This grouping is also reflected vertically in the top annotation track, where each sample is labeled by its biological condition. The genes (rows) are clustered across samples based on Z-scores for expression, resulting in scaling genes (pinpointing variability found within genes [33].

**Figure 7:**
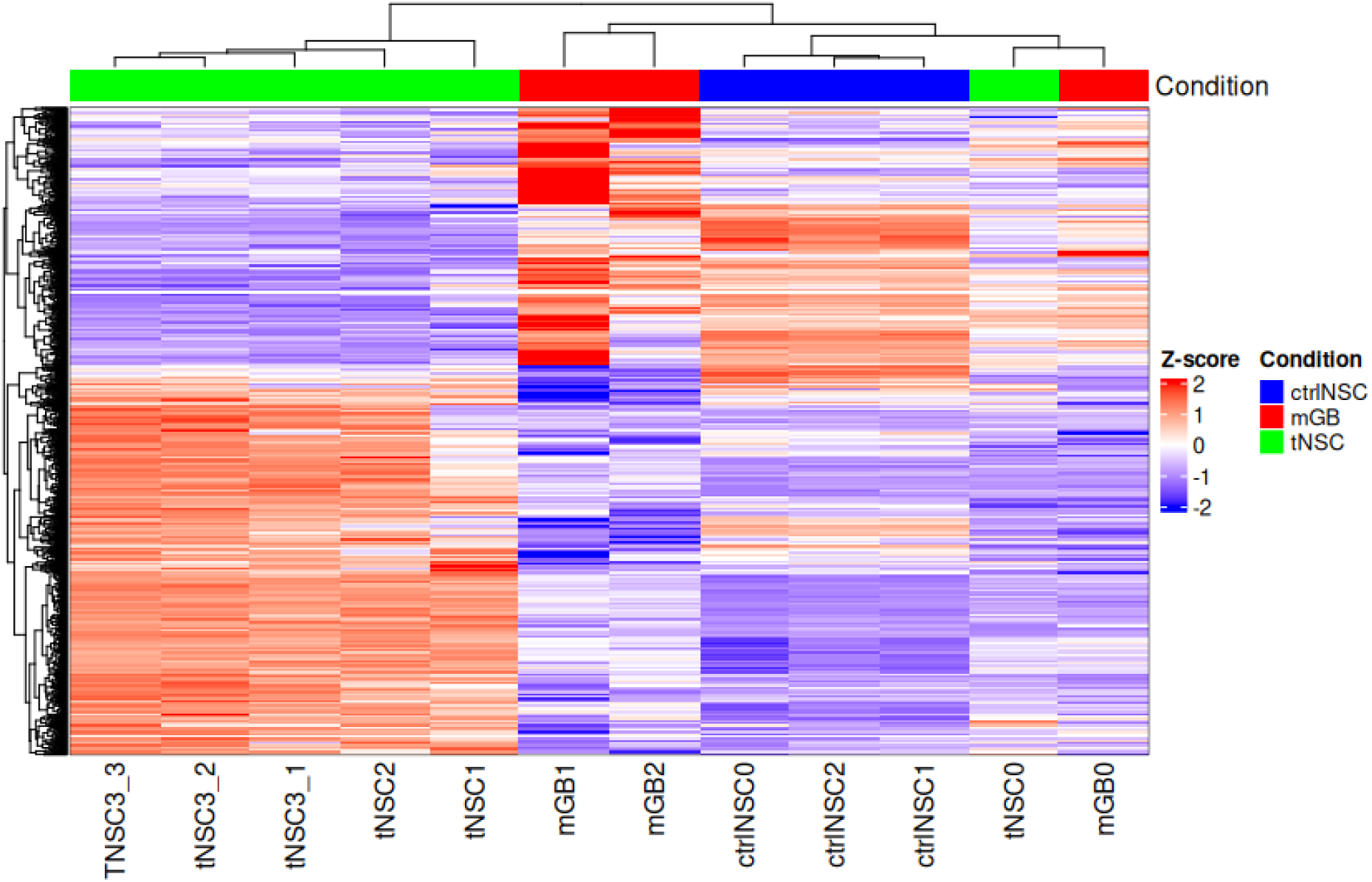
Heatmap of differential expression across all the profiles.

The tNSC samples (tNSC0–tNSC3) shared an expression profile that is between the extremes of mGB and ctrlNSC, suggesting that tNSC has a molecular phenotype in transition. Of note, mGB1 displayed a distinctly different pattern of expression with strong upregulation (represented as red) and downregulation (represented as blue) of specific gene clusters, reflecting either the aggressive tumor phenotype of mGB1 or temporal (state) heterogeneity. In contrast, ctrlNSC samples clustered closely together, suggesting that in non-tumor cells the transcriptional program is stable and consistent.

This global heat map highlights transcriptional reprogramming that occurs during gliomagenesis. Differential expressed gene clusters (located in the middle of the heat map) are either activated or repressed specifically in mGB sample, as opposed to both tNSC and ctrlNSC, suggesting involvement in tumor-specific regulatory pathways.

## 4 Discussion

This study focuses on whole-exome sequencing (WES), Ti/Tv profiling, mutational signature analysis, Gene Ontology and transcriptomic data to investigate the molecular processes underlying GBM progression in TP53-PTEN deficient murine model. These analyses revealed a two-phase evolutionary trajectory: an early phase where samples demonstrated genomic instability and developmental pathway disruption in transformed neural stem cells (tNSCs), and a later phase where oncogenic refinement, transcriptional plasticity, and invasive phenotypes were characterized called malignant glioblastoma (mGBs). Below we explain the molecular interplay and transcriptional dynamics across these distinct stages.

### 4.1 Mutational Evolution: From Genomic Chaos to Oncogenic Convergence

#### 4.1.1 Early-Stage Genomic Instability

The tNSC phase exhibited a high burden of transversion-biased mutations (Ti/Tv = 1.18-1.66), consistent with the mutational signatures SBS4 (oxidative stress) and SBS40 (developmental processes) [28]. These are often assiciated with the loss of TP53 and PTEN, imparing DNA repair and amplifies oxidative damage, crafting suitable environment for early mutagenesis. SBS26 (mismatch repair deficiency) dominance across all samples indicated MMR deficiency as initiating event, causing mutation accumulation. This contrasts with human GBM, where MMR deficieny is often observed especially post-treatment. Moreover, Pfdn2, Oip5, Trmt10a, Man2b1, and Trem1 lays the foundational mutations in transforming neural stem cells to malignancy.

#### 4.1.2 Late-Stage Oncogenic Refinement

In mGBs, Ti/Tv ratios increased to 1.96-2.02, implicating a shift to replication-associated transitions (SBS5/SBS37) and APOBEC-like mutagenesis. This transition also parallels human GBM progression, where advanced tumors acquire SBS5 and oncogenic drivers under selective pressure. Importantly, recurrent mutations in Ubap2, Tead2, Errfi1, Apobec2 and CDK4 (cell cycle) were found only in mGBs, supporting the notion of a hierarchial model in which genome chaos early on in tumor progression, promoted pathway convergence later on. The lack of these mutations in tNSCs supports a role in malignant transformation vs initiation.

### 4.2 Functional Impact: Linking Mutations to Biological Processes

#### 4.2.1 GO Analysis of High/Moderate-Impact Variants

The early tNSCs were enriched for neurodevelopment pathways (neurogenesis, gliogenesis), driven by mutations in CTNNB1 (Wnt-*β*-catenin) and NF1 (RAS/MAPK) [30]. These alterations appears to disrupt stem cells differentiation, locking tNSCs in a proloferative, differentiated state. In mGBs, Gene Ontology terms shifted towards cell junction assembly and microtubule-based movement. These changes coincides with mutations in CDK4 that causes invasion and cell cycle dysregulation. This molecular divergence reflects transitions causing development destabilization to structural remodeling (a hallmark of mesenchymal GBM).

#### 4.2.2 Transcriptional Plasticity Mirrors Tumor Evolution

‘The transcriptomic landscape of GBM progression, as revealed by principle component analysis (PCA) and hierarchial clustering, undermines the transcriptional reprogramming that distinguishes normal neural stem cells (ctrlNSC), pre-malignant (tNSC) and malignant glioblastoma (mGB). This difference can be attributed to the activation of oncogenic pathways such as cell cycle regulators and migratory genes, but also the repression of neural differentiation programs, as observed with tNSCs occupying a position intermediate of ctrlNSCs and mGBs. The separation of mGBs along PC2 (11% variance) means they have become transcriptionally unique because they consistently have increased proliferation (Cdk4, Mki67) and invasion-associated gene transcription. These transcriptional “states” are concurrent with the somatic mutations seen through WES analysis for the Bradford cohort, with late stage CDK4 mutations corresponding with transcriptional pathway activation, suggesting a genotype-to-phenotype relationship.

#### 4.2.3 Heterogeneity in Pre-Malignant States

The intra-group variance noted in tNSCs (showing dispersed clustering in PCA) indicates heterogeneity early in gliomagenesis. This could be due to arbitrary epigenetic reprogramming, different exposures to tumor microenvironmental factors, or asynchronous progression in tNSC subclones. In fact, tNSC1 and tNSC2’s proximity to mGBs likely indicates they have progressed to advanced pre-malignant states and are on the edge of transformation. Meanwhile, tNSC3’s closeness to ctrlNSCs could indicate it has held onto stemness or just has a delayed commitment to become malignant. This type of heterogeneity is similar to human GBM, which is clinically important, as intratumoral heterogeneity complicates the targeting of therapeutics and drives recurrence.

### 4.3 Malignant Transition and Regulatory Pathways

The clustered heatmap of the transcriptome demonstrates three transcriptional groups (ctrlNSC, tNSC, mGB) with mGBs showing clear and significant upregulation of gene clusters that are specific to tumors. For example, the high overexpression of Cdk4 in mGBs relates to its recurrent somatic mutations and supports its function in cell-cycle dysregulation. The clusters downregulated in mGBs include genes for neural differentiation and genes involved in synaptic signaling and represent a loss of stemness and gain of among-mesenchymal characteristics. The transcriptomic clusters also support the GO enrichment analysis of somatic variants, when late-stage mutations associated with structural and motility pathways (i.e. cell junction assembly, microtubule-based movement) become dysregulated to drive transcriptional reprogramming to invasive phenotypes.

### 4.4 Clinical Implications and Future Directions

The transcriptional differences between tNSCs and mGBs present potential opportunities:

**Key therapeutic target:** Cdk4 and other upregulated genes in mGB provide reasoning for treating patients with CDK4/6 inhibitors (e.g., ribociclib) along with radiotherapy, an approach that is actively being evaluated in clinical trials.

**Heterogeneity considerations:** the diversity that markdowns tNSCs, demonstrate the necessity for future single-cell transcriptomic studies so we can deconvolve subclonal populations and identify early event drivers of transformation.

#### Limitations

This work demonstrates important transcriptional trends, but without longitudinal sampling we cannot map real-time evolution. Additionally, the murine model we used is not representative of immune and stromal interactions, which could diminish the extent to which the microenvironment will affect transcriptional plasticity.

